# Limited genetic parallels underlie convergent evolution of quantitative pattern variation in mimetic butterflies

**DOI:** 10.1101/2020.06.15.151613

**Authors:** Hannah E. Bainbridge, Melanie N. Brien, Carlos Morochz, Patricio A. Salazar, Pasi Rastas, Nicola J. Nadeau

## Abstract

Mimetic systems allow us to address the question of whether the same genes control similar phenotypes in different species. Although widespread parallels have been found for major effect loci, much less is known about genes that control quantitative trait variation. In this study, we identify and compare the loci that control subtle changes in the size and shape of forewing pattern elements in two *Heliconius* butterfly co-mimics. We use quantitative trait locus (QTL) analysis with a multivariate phenotyping approach to map the variation in red pattern elements across the whole forewing surface of *Heliconius erato* and *Heliconius melpomene*. These results are compared to a QTL analysis of univariate trait changes, and show that our resolution for identifying small effect loci is improved with the multivariate approach. QTL likely corresponding to the known patterning gene *optix* were found in both species but otherwise, a remarkably low level of genetic parallelism was found. This lack of similarity indicates that the genetic basis of convergent traits may not be as predictable as assumed from studies that focus solely on Mendelian traits.

## Introduction

Evolutionary convergence is the independent evolution of the same phenotype in response to the same ecological challenge (Stern 2013). Convergent systems allow us to ask whether similar selection pressures result in similar genetic changes (Hoekstra and Nachman 2003). Some researchers hypothesise that homologous loci (*i.e.* the same genes) will be repeatedly used by unrelated species to produce identical phenotypes, a process called ‘parallel genetic adaptation’ (Wood et al. 2005; Joron et al. 2006; Stern 2013). Finding strong parallels at the genetic level suggests that there are genetic, developmental or evolutionary factors which make certain genes more likely targets of natural selection. Nevertheless, there are also some clear cases where convergence results from different genes (Hoekstra and Nachman 2003; Ng et al. 2008). For example, Roelants et al. (2010) report a striking example of convergence between two frog lineages, *Xenopus* and *Litoria*, which independently evolved the same toxic skin secretions via different precursor genes (*cholecystokinin* and *gastrin*, respectively). Therefore, the extent to which phenotypic convergence reflects similar or divergent genetic mechanisms is still being debated (Stern 2013).

Theoretical work suggests that adaptive evolution likely follows a ‘two-step’ model, where species initially undergo rapid phenotypic change, controlled by large effect loci, followed by smaller changes in order to reach a final trait optimum (Nicholson 1927; Sheppard et al. 1985; Orr 1998; Baxter et al. 2009). It is considerably easier to identify loci of large effect than those of small effect which control subtle variation (Beavis 1998; Rockman 2011). Although high levels of genetic parallelism are often found at major effect loci between convergent species, it is likely these findings do not represent the true complexity of the genetic architecture underlying adaptive traits (Shapiro et al. 2004; Colosimo et al. 2005; Joron et al. 2006; Kronforst et al. 2006; Papa et al. 2008; Morris et al. 2019). Thus, if we move the focus of research away from discrete traits and towards the more subtle variation within quantitative traits, a host of small effect loci could be revealed, that may differ in the extent of parallelism they exhibit (Wood et al. 2005).

*Heliconius* butterflies have undergone adaptive radiations into multiple colour pattern races across the neotropics, with widespread phenotypic convergence between both closely and distantly related species, making the genus an ideal study system for investigating the level of genetic parallelism that underlies repeated evolution (Papa et al. 2008; Hines et al. 2011; Nadeau et al. 2014). These toxic, warningly coloured species converge onto the same wing colour patterns as a mechanism to enhance predator avoidance learning, a process known as Müllerian mimicry (Benson 1972; Sheppard et al. 1985). In turn, the resultant shared local mimicry between species allows us to ask if parallel patterns of divergence in different species are generated using parallel genetic mechanisms (Parchem et al. 2007; Nadeau and Jiggins 2010).

To date, most of our knowledge of the genetic control of wing pattern variation in *Heliconius* comes from studies focusing on major Mendelian genes, which control specific pattern elements (Papa et al. 2008). These studies have identified a set of 5 ‘tool-kit’ loci, thought to control most of the pattern variation across the genus (Sheppard et al. 1985; Naisbit et al. 2003; Joron et al. 2006; Kronforst et al. 2006; Baxter et al. 2009; Hines et al. 2011; Papa et al. 2013; Nadeau 2016). *Heliconius* has therefore been one of the taxa contributing to the paradigm that a high level of genetic parallelism exists between convergent species. However, there is limited genetic information for the more subtle variation in colour patterns, which is likely to be controlled by a larger number of small effect loci (Fisher 1930; Wood et al. 2005). The handful of studies that have investigated quantitative pattern variation in *Heliconius* found that the tool-kit loci also control this subtler variation in pattern to a greater or lesser extent (Baxter et al. 2009; Jones et al. 2012; Papa et al. 2013; Huber et al. 2015; Van Belleghem et al. 2017; Morris et al. 2019). However, no studies have directly investigated the extent of parallelism in the loci and architecture of quantitative trait variation in co-mimetic races.

In this study, we use the program *patternize* (Van Belleghem et al. 2018) to extract quantitative variation in red forewing (FW) pattern between the co-mimics *Heliconius melpomene* and *Heliconius erato*. Such analyses can objectively capture subtleties in pattern variation across the entire wing surface, allowing us to consider multiple axes of trait variation at once (hereafter, multivariate trait analysis), rather than simple discrete trait changes (Jones et al. 2012; Huber et al. 2015). The power of mapping studies to detect causal loci has been shown to greatly increase when the multivariate nature of trait changes are considered (Schmitz et al. 1998). Therefore, we use this approach to address the question of whether the same loci with the same distribution of effect sizes have been used during the convergence of these two species.

## Methods

### Experimental crosses

*H. erato demophoon* and *H. melpomene rosina* individuals from Gamboa, Panama (9.12°N, 79.67°W) were single pair mated with *H. erato cyrbia* and *H. melpomene cythera* individuals, respectively, from Mashpi, Ecuador (0.17°N, 78.87°W), to produce F_1_ hybrid offspring. Crossing F1 individuals together then generated F2 offspring, whilst one *H. erato* backcross (BC) was produced by mating an F1 male with a *H. e. cyrbia* female (Figure 1). The *H. erato* broods were used in a previous analysis of iridescent colour variation (Brien et al. 2018). Within a few hours of emergence, adult wings were removed and stored in glassine envelopes and bodies were stored in NaCl saturated 20% dimethyl sulfoxide (DMSO) 0.25M EDTA solution to preserve the DNA. In total, 3 *H. melpomene* families were sequenced and used in the final analyses (2 grandparents, 5 parents and 113 F2 offspring), along with 6 *H. erato* families (5 grandparents, 11 parents, 99 F2 offspring and 40 backcross offspring). A further 184 *H. melpomene* and 93 *H. erato* individuals were included in the PCA analysis. Full details of all crosses are in Table S1.

**Figure 1.**
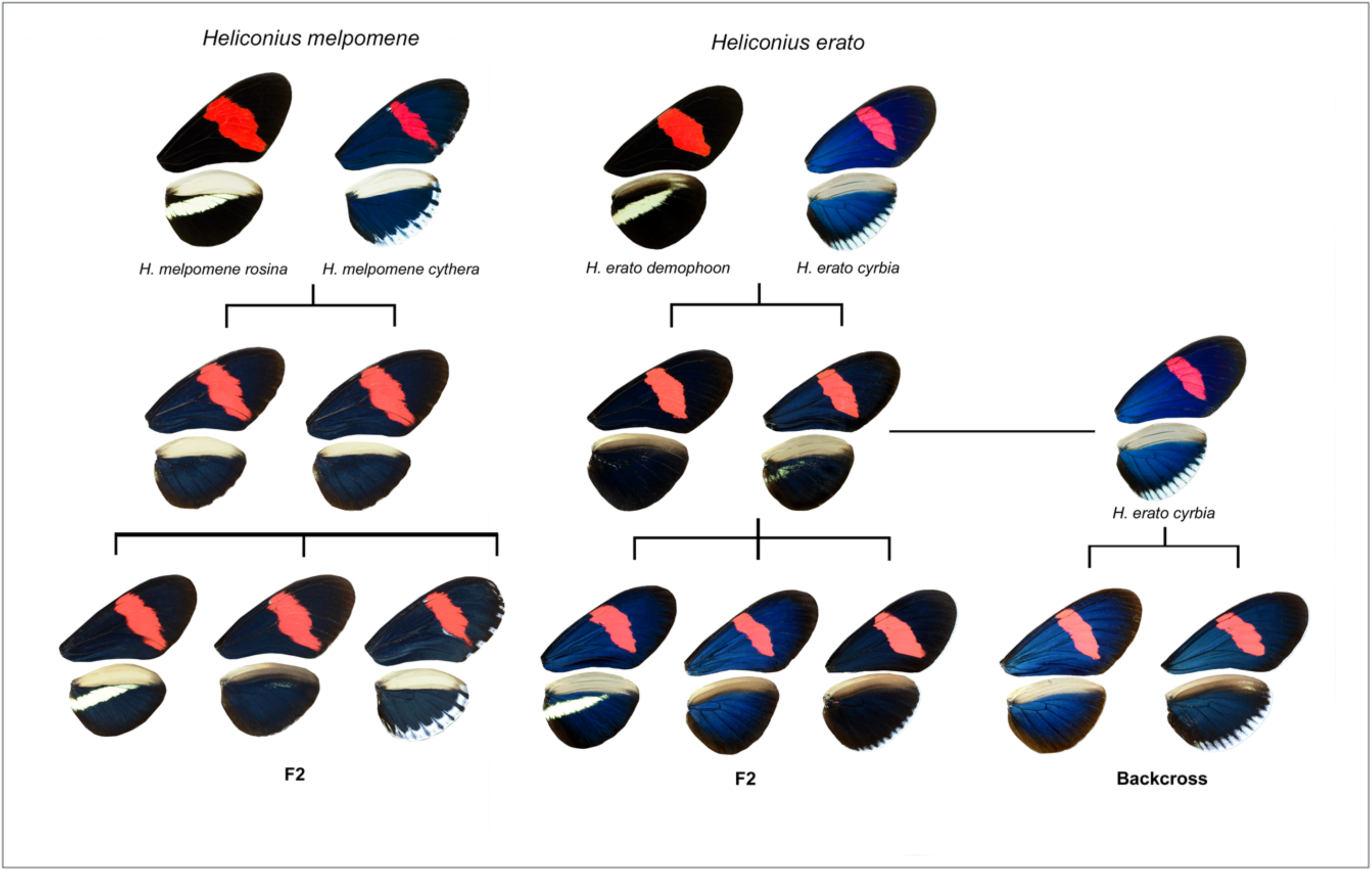
Cross-design and examples of colour pattern variation in *H. melpomene* and *H. erato*: F1, F2 and backcross (BC) generations (adapted from Brien et al. 2018).

### DNA extraction and sequencing

Genomic DNA was extracted using the Qiagen DNeasy Blood & Tissue kit following the manufacturer’s instructions, with an additional treatment with Qiagen RNase A to remove RNA. Approximately half of the thorax of each individual was used in the extraction. Single-digest Restriction site-associated DNA (RAD) library preparation and sequencing were carried out by the Edinburgh Genomics facility at the University of Edinburgh. DNA was digested with the enzyme *PstI*, which has a cut site approximately every 10kb. Libraries were sequenced on an Illumina HiSeq 2500 producing 125bp paired-end reads. Available parents of the crosses were included at 2x higher concentration within the pooled library to produce a higher depth of coverage.

### Sequence data processing

The RADpools function in RADtools version 1.2.4 was used to demultiplex the RAD sequences, using the option to allow one mismatch per barcode (Baxter et al. 2011). Quality of all raw sequence reads were checked using FastQC (version 0.11.5, Babraham Bioinformatics). FASTQ files were mapped to the *H. erato* v1 reference genome (Van Belleghem et al. 2017) or the *H. melpomene* v2.5 reference genome (Davey et al. 2017) using bowtie2 v2.3.2 (Langmead and Salzberg 2012). BAM files were then sorted and indexed with SAMtools (v1.3.1). PCR duplicates were removed using Picard tools MarkDuplicates (v1.102). Genotype posteriors were called with SAMtools mpileup (Li 2011), set to a minimum mapping quality of 10 and minimum base quality of 10, using Lep-MAP data processing scripts (Rastas 2017).

### Genetic map construction

Linkage maps were constructed using Lep-MAP3 (Rastas 2017). Before starting, the sex of each individual was confirmed by comparing the depth of coverage of the Z chromosome against a single autosome. Females have half the depth of coverage on the Z compared to the autosomes, as they only have one copy of the Z. Based on this, one *H. melpomene* individual was removed when the inferred sex could not be verified using the wings. The IBD module was run to verify the pedigree and a further 2 individuals (1 *H. erato*, 1 *H. melpomene*) were removed when the ancestry could not be confirmed.

The ParentCall2 module was used (with options ZLimit=2, removeNonInformative=1) to calculate and impute the most accurate parental genotypes and sex chromosomal markers using information from related parents, grandparents and offspring provided as a pedigree file. Markers were then filtered to remove those with high segregation distortion (dataTolerance=0.001). Next, markers were assigned to 21 linkage groups with the SeparateChromosomes2 module which calculates LOD scores between pairs of markers. For *H. melpomene*, LOD limits between 10 and 25 were tested within this module, and a limit of 23 used as this gave the correct number of linkage groups with an even distribution of markers. For *H. erato*, markers were separated into 21 linkage groups based on the chromosomal reference genome, then a LOD limit of 10 was used only between markers in each of these groups, keeping the largest group per chromosome.

JoinSingles2all added additional single markers to the existing linkage groups. OrderMarkers2 was used to order the markers within each linkage group by maximising the likelihood of the data. In this, recombination2 was set to 0, because there is no recombination in females (Suomalainen et al. 1973) and male recombination set to 0.05 (following Morris et al. 2019). Using parameter outputPhasedData=1 gave phased data and imputed missing genotypes and, for the *H. melpomene* map, additional parameter hyperPhaser was provided to improve phasing of markers. For the *H. erato* map, the marker order was further evaluated via parameters evaluateOrder=order.txt and improveOrder=0, where order.txt contained the markers in the physical order.

Finally, we used LMPlot to visualise the maps and check for errors in marker order. Any erroneous markers that caused long gaps at the beginning or ends of the linkage groups were manually removed. Genotypes were phased using Lep-MAP’s map2genotypes.awk script, and markers were named using the map.awk script and a list of the SNPs used to provide the scaffold name and position. Markers which had grouped to the wrong linkage group based on genomic position were removed (<1% of the total markers). The final genetic map for *H. erato* contained 5648 markers spread over 21 linkage groups with a total length of 1162.4cM (Table S2), and for *H. melpomene* the final map had 2163 markers spanning 1469.9cM (Table S3).

### Phenotypic analysis of the broods

Wings were photographed using a mounted Nikon D7000 DSLR camera with a 40mm f/2.8 lens set to an aperture of f/10, shutter speed of 1/60 and ISO of 100, and paired Kaiser Fototechnik RB 5004 lights with daylight bulbs. All photographs also included an X-Rite Colour Checker to help standardise the colour of the images. RAW format images were standardised using the levels tool in Adobe Photoshop CS2 (Version 9.0).

Red, green and blue (RGB) values for red pattern elements were selected via the *patternize* sampleRGB() function in R (R Core Team 2018; Van Belleghem et al. 2018). These were 218, 77, 46 for *H. erato* and 211, 1, 30 for *H. melpomene. patternize* used these values to extract the presence and distribution of red pattern elements across the dorsal side of each forewing. ‘Offset’ values were incrementally increased until coloured areas included as much of the red forewing pigmentation as possible, whilst minimising the amount of background ‘noise’ included (Hemingson et al. 2019). The final values were 0.24 for *H. erato* and 0.35 for *H. melpomene.*

To align the images, eighteen landmarks at wing vein intersections (Figure S1) were manually set, using the Fiji distribution of ImageJ (Schindelin et al. 2012). Landmark registration within the *patternize* package was used to align and extract the red pattern elements from each generation of the crosses. The variation extracted was summarised in a principal component analysis (PCA), allowing us to examine the patterns of variation and co-variation both among the red pattern element data and across the different generations (Lee et al. 2018). Additionally, patArea() was used to calculate the relative area of the colour pattern.

Linear measurements were also taken on the dorsal side of the forewings, to measure how far towards the distal edge of the forewing the red band extended (hereafter, ‘distal extension of the red FW band’). Using wing veins as fixed reference points, a measurement was taken along the lower edge of the red FW band. Three other wing measurements were also taken to standardise for wing size (Baxter et al. 2009) (Figure S2). Measurements were carried out using ImageJ and repeated for both left and right wings. Final values were then calculated by taking the average of the two band measurements and dividing it by the average of all the standardising measurements.

### Quantitative Trait Locus mapping analysis

QTL scans were carried out across the 20 autosomes using R/shapeQTL (Navarro 2015) for multivariate traits (principal components from the PCA) and R/qtl (Broman and Sen 2009) for univariate traits (red band area and linear measurement). Analyses were run separately for *H. melpomene* F2, *H. erato* F2 and the *H. erato* backcross. The Z sex chromosome had to be excluded from the R/shapeQTL analysis but was included in the R/qtl analysis. Sex was included as an additive covariate in all analyses to account for possible sexual dimorphism in FW band elements (Klein and de Araújo 2013). For analysis of F2 crosses, family was also included as a covariate.

For R/qtl analyses, the Haley-Knott (HK) algorithm was implemented to map QTL for area and FW band extension. Statistically significant LOD thresholds were calculated using 1000 permutations (Churchill and Doerge 1994) with *perm.Xsp=T* to get a separate threshold for the Z chromosome. The test for linkage on the sex chromosome has 3 degrees of freedom compared to 2 for the autosomes, so more permutation replicates are needed (21832 for *H. erato* and 12918 *H. melpomene*, compared to 1000 permutations on autosomes) and the significance threshold is higher for the Z chromosome (Broman et al. 2006). For each QTL above the LOD threshold, 95% Bayesian credible intervals were computed to refine its boundaries. Effect sizes of the significant QTLs were then estimated using fitqtl().

For R/shapeQTL analyses, a multivariate Pillai trace test was used to map traits. Here, it is necessary to limit the number of variables (PCs) relative to the number of samples in order to make the estimates more robust (Maga et al. 2015; Morris et al. 2019). In these analyses, all PCs explaining >1% of the variation were included. Eight PCs (each explaining 10%, 5.5%, 2.8%, 1.8%, 1.7%, 1.4%, 1.2% and 1% of the variation, respectively) were used in the *H. melpomene* QTL analysis, whilst 9 PCs were included in the *H. erato* QTL analysis for both F2 and backcrosses (explaining 9.7%, 4.5%, 3.5%, 2.1%, 1.9%, 1.4%, 1.3%, 1.2 % and 1% of the variation) (Figure S3). Cumulatively, these explained only 25.4% of the overall variation in *H. melpomene* and 26.6% in *H. erato*.

We used LepBase (Challis et al. 2016) to locate the nearest gene and its Gene Ontology (GO) molecular function for markers with the highest LOD scores in significant QTL. If any previously identified *Heliconius* wing patterning loci were known to fall on a chromosome containing a QTL, we determined whether these fell within the 95% Bayesian credible intervals of these QTL. The homologue of the *Drosophila ventral veins lacking (vvl)* gene (required for cell proliferation and differentiation of wing veins (Sturtevant et al. 1994)) was treated as the patterning gene on chromosome 13, which has been identified in previous mapping studies (Van Belleghem et al. 2017; Morris et al. 2019), but not functionally verified.

To test for broad-scale genetic parallelism between species and to look for additive effects of multiple small effect loci, we determined the effect sizes of whole chromosomes by running a genome scan in the F2 crosses for both the univariate and multivariate traits using only maternal alleles. As there is no recombination of maternal chromosomes, maternal markers can be used to investigate chromosome level effects. To do this, the effects of paternal markers were ignored by setting all paternal markers to be the same allele (here we used ‘A’). The *H. erato* and *H. melpomene* genomes are known to be highly syntenic (Kronforst et al. 2006; The Heliconius Genome Consortium 2012; Van Belleghem et al. 2017), therefore, to test for parallelism we assessed whether there was a correlation between these chromosome-level effects between species, using a Spearman’s Rank correlation.

## Results

### Multivariate analysis of red band shape

Heat maps were used to visualise variation in forewing band shape and compare this between *H. erato* and *H. melpomene*. Interestingly, although the heat maps for the F2 generation showed that the two species exhibited similar variation, the shape of the FW bands for *H. melpomene* and *H. erato* appears to be slightly different. *H. melpomene* showed marginally less overall variation and possessed a FW band that extended further towards the distal edge. However, it is clear that for both species, most of the variation occurred around the border of the FW band (Figure 2 A, B). The parental races were non-overlapping in phenotypic space in the PCA. As expected for a genetically controlled trait, the F1 offspring were intermediate, with F2 offspring having a wider distribution of phenotypic values that overlap with those of the parental races, and backcross offspring being skewed towards the backcross parent (Figure 2 C, D).

**Figure 2.**
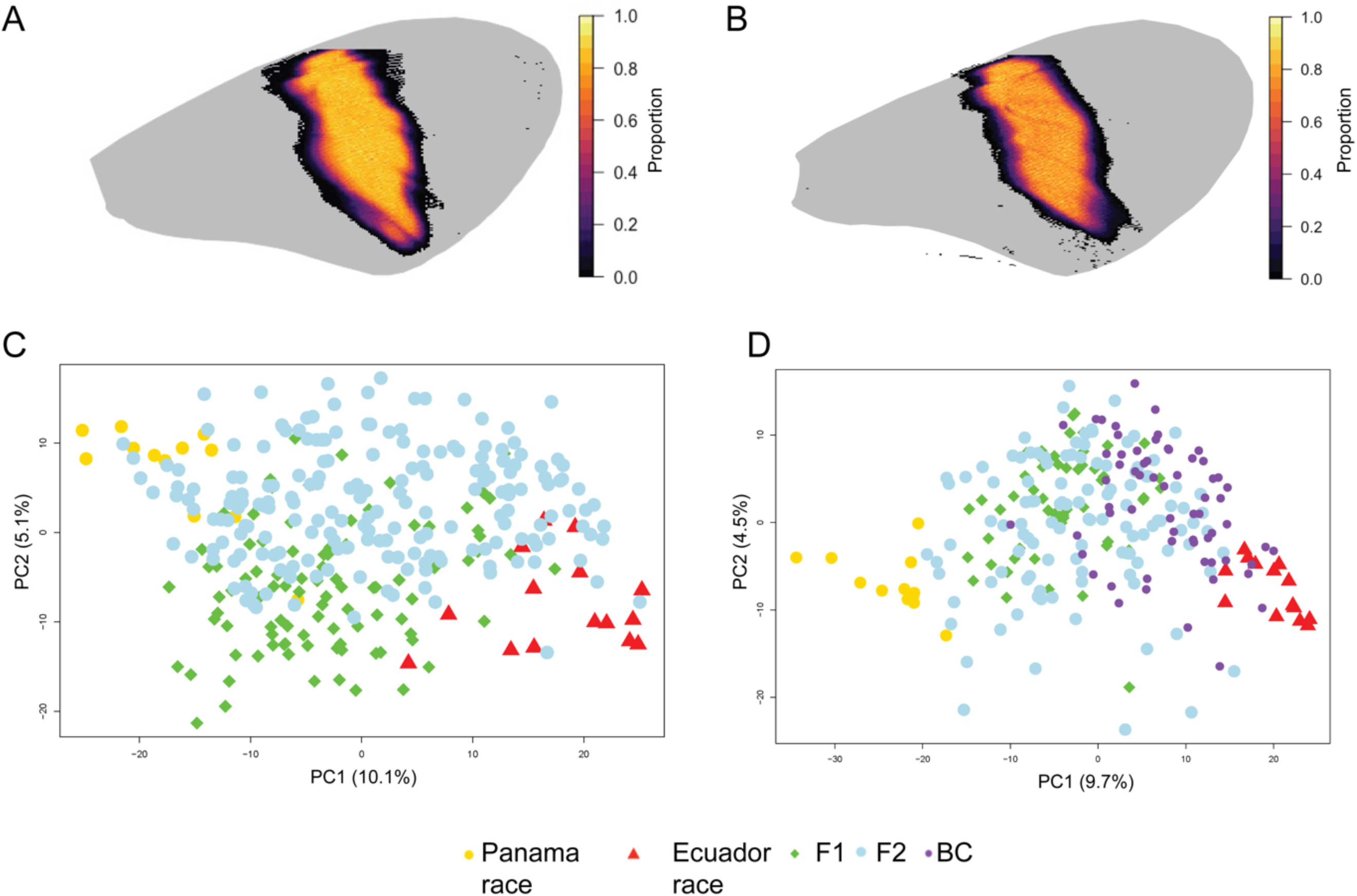
Multivariate analysis of quantitative red FW variation, showing heat maps for the variation in the F2 generation of *H. melpomene* (A) and *H. erato* (B). The colour relates to the proportion of individuals that have red pigmentation at each pixel, where darker colours indicate greater variation. Principal component analyses of all individuals across the different crossing generations show how the parental variation segregates in *H. melpomene* (C) and *H. erato* (D). Note only PC1 and 2 are shown here.

To find genomic regions controlling the variation captured by the PCA, QTL scans were conducted for the F2 generations of both species and for the *H. erato* BC family. No LOD scores were above the significance threshold for the BC generation. However, in the *H. erato* F2 generation, three QTL were found on chromosomes 11, 13 and 18. These explained 3.4%, 4.6% and 5.9% of the variation in FW red colour pattern, respectively. For *H. melpomene*, a single significant QTL was found, also on chromosome 18. This explained 4.3% of the variation (Figure 3 A, B and Table 1). Chromosomes 13 and 18 harbour the known ‘tool-kit’ loci *Ro* (*vvl*) and *optix* respectively. In both species, the 95% Bayesian credible intervals on chromosome 18 overlapped with the gene *optix* (Figure 4), although the interval was large in *H. melpomene*, spanning almost the entire chromosome. In the *H. erato* map, the marker with the highest LOD score was only 1.5cM from the position of *optix*. The 95% Bayesian credible interval of the QTL on *H. erato* chromosome 13, however, did not contain the gene *vvl*.

**Table 1.**
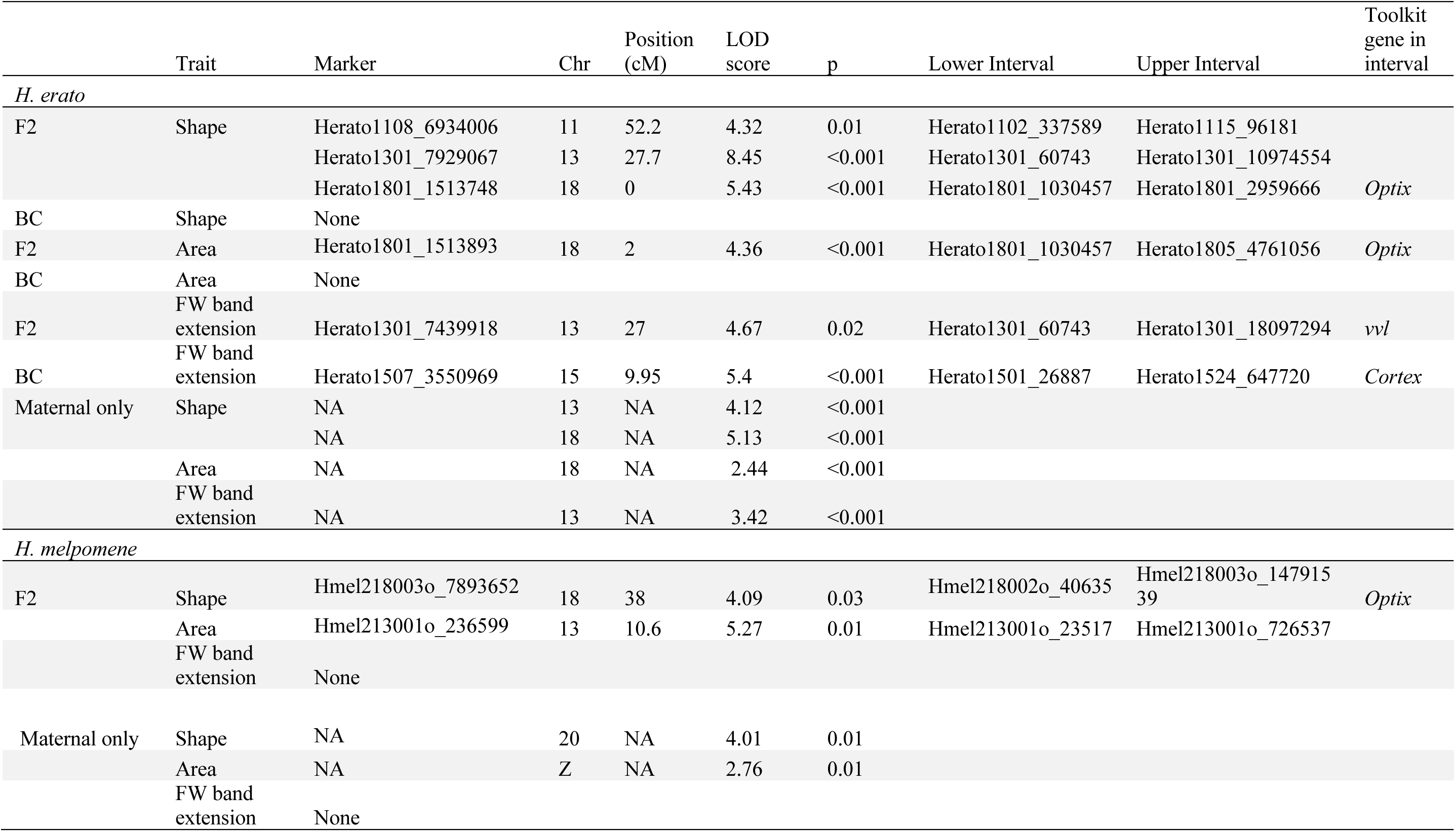
All significant QTL found in the multivariate analyses for shape and univariate analyses for area and FW band extension. Tool-kit loci are noted where they occur within the 95% Bayesian intervals of the QTL. Details of tool-kit loci positions within the reference genomes can be found in Table S4.

**Figure 3.**
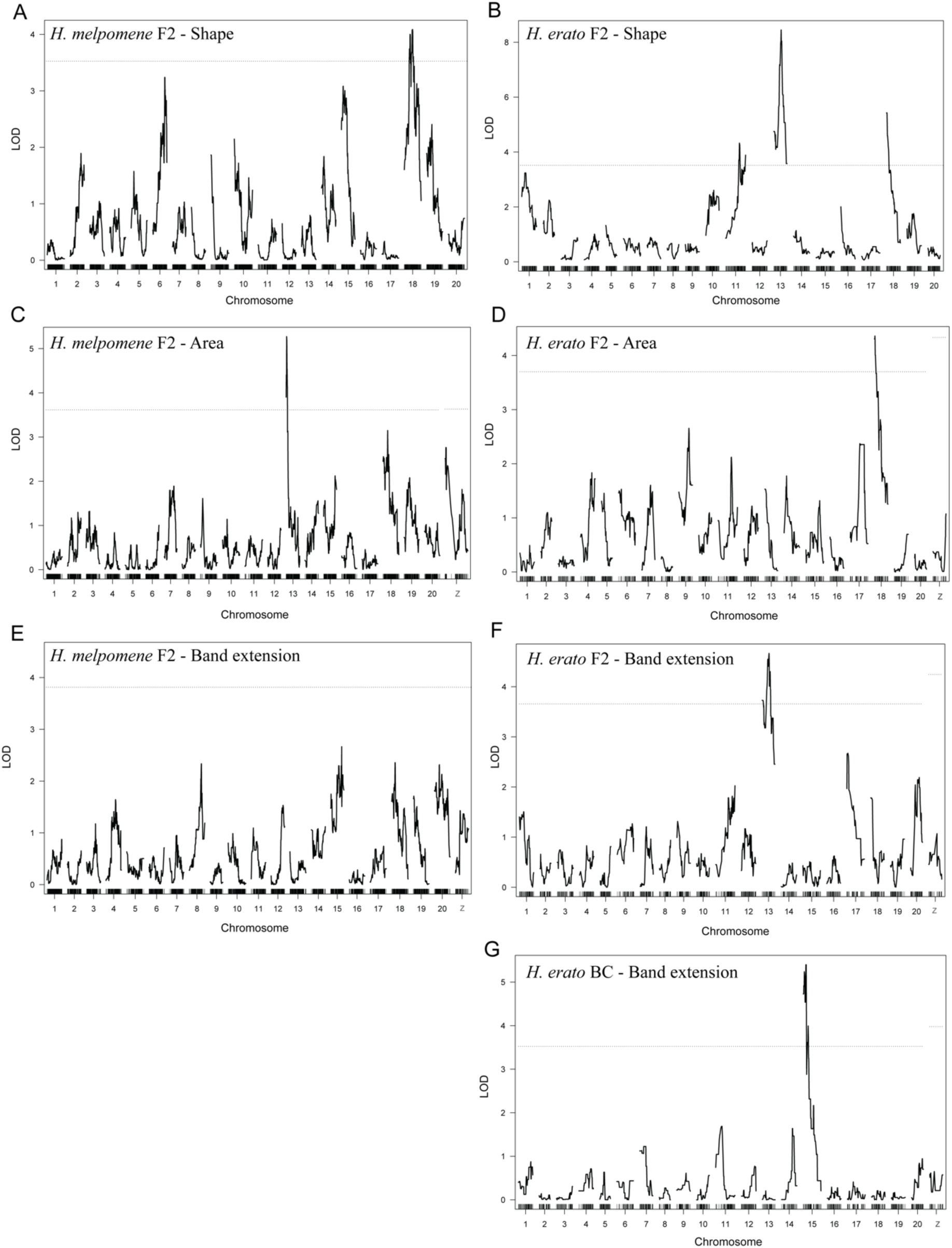
ShapeQTL scans of the F2 generation of *H. melpomene* (A) and *H. erato* (B) for the red pattern variation (captured by the PCA). Dotted lines show significance thresholds for additive LOD scores. QTL scans for relative area of the red forewing bar in F2 *H. melpomene* (C) and *H. erato* (D). QTL scans for extension of the forewing band in *H. melpomene* (E), *H. erato* F2 (F), and *H. erato* backcross (G) generations. There were no significant QTL for band extension found in *H. melpomene*.

**Figure 4.**
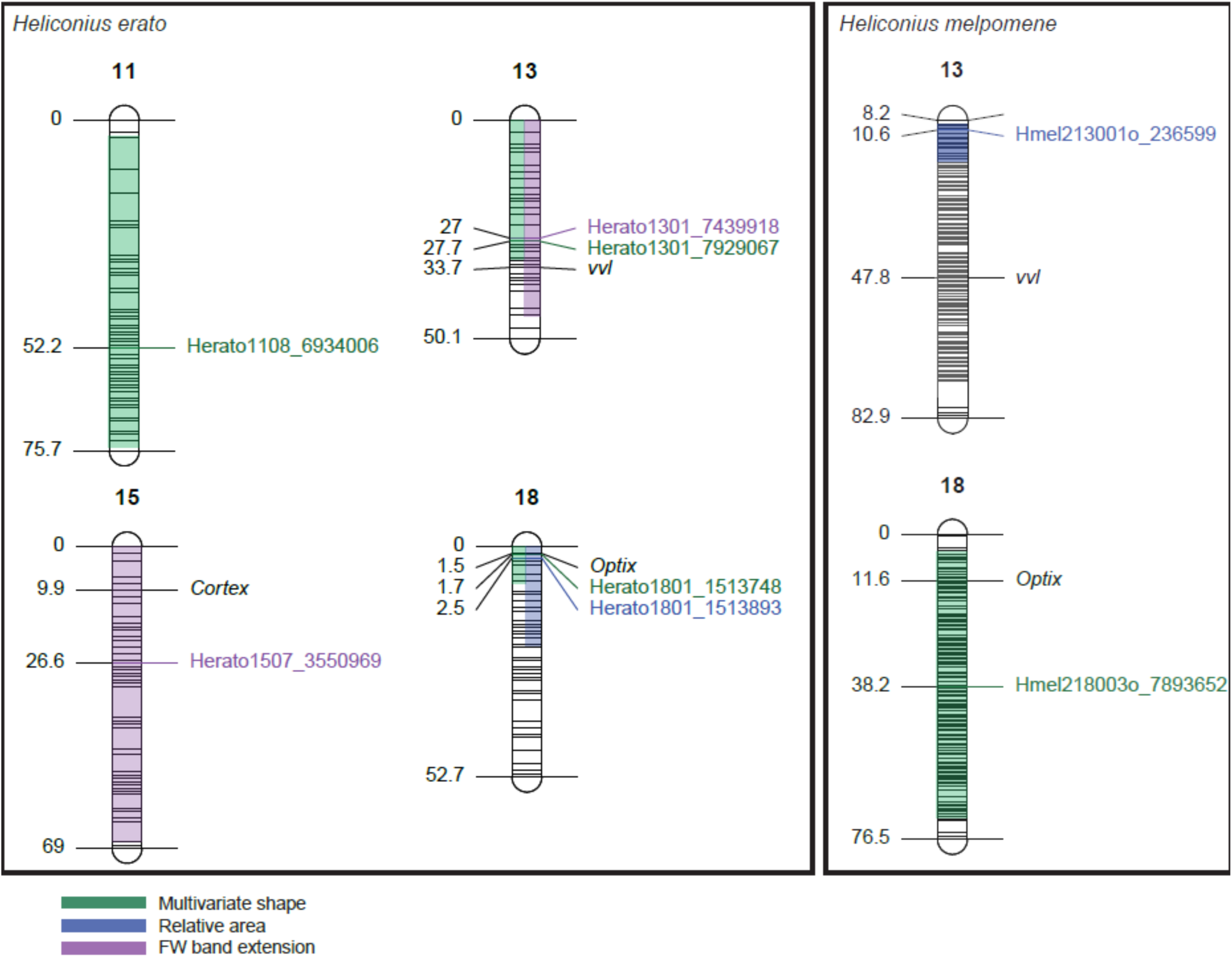
An overview of the locations of the QTL found in the analyses for multivariate shape (highlighted in green), relative area (blue) and FW distal band extension (purple). 95% Bayesian credible intervals are highlighted in the respective colours. Position on the linkage group (cM) is shown on the left of each chromosome, and the name of the significant markers are shown on the right. Positions of the toolkit loci (o*ptix, cortex* and *vvl*) are also shown. Figure created using LinkageMapView (Ouellette et al. 2018).

### Univariate analysis of relative area of the FW band

We also performed QTL mapping of the univariate trait, relative area of the red FW band, in both species. As with the multivariate trait analysis, no LOD scores were found above the significance threshold for the backcross generation. However, for the F2 generations of both *H. melpomene* and *H. erato*, single significant QTL were found (Figure 3 C, D). In *H. melpomene*, this was on chromosome 13 (LOD = 5.27, p = 0.01) and explained 17.4% of the variation, and in *H. erato* the QTL was on chromosome 18 (LOD = 4.36, p<0.001) and explained 18.9% of the variation. *Optix* was within the 95% Bayesian credible intervals for the *H. erato* chromosome 18 QTL, and thus could be a candidate locus here (Figure 4, Table 1). The gene *vvl* was not within the 95% Bayesian credible interval for the *H. melpomene* chromosome 13 QTL.

Effect plots show that in *H. erato*, heterozygotes for the chromosome 18 marker were on average closer in phenotype to homozygotes for the *demophoon* (the Panama subspecies with large red bar) allele than homozygotes for the *cyrbia* allele, suggesting slight dominance of *demophoon*, consistent with the general pattern whereby the presence of red pattern elements controlled by *optix* tends to be dominant. In contrast, in *H. melpomene*, heterozygotes for the chromosome 13 marker expressed slightly smaller FW bands than either homozygote, sometimes referred to as ‘underdominance’ (Figure 5 A, B).

**Figure 5.**
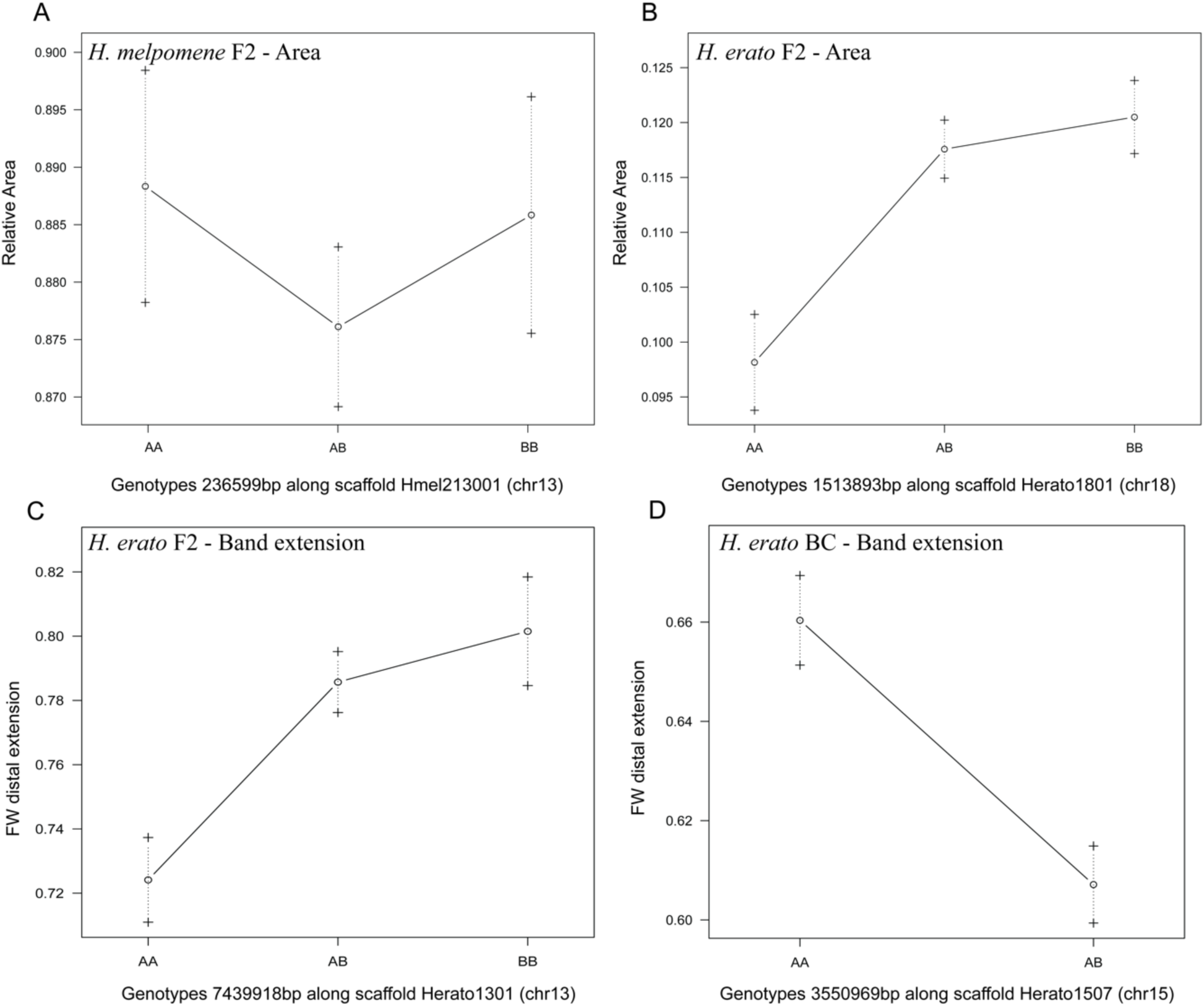
Effect plots showing the FW band area phenotype means for each group in the F2 generation of *H. melpomene* (A) and *H. erato* (B), defined by the genotypes at the respective markers. Effect plots showing the phenotype means for FW band extension in the *H. erato* F2 (C) and BC generations (D). The ‘A’ alleles are from the iridescent race (*H. m. cythera* and *H. e. cyrbia*) and the ‘B’ alleles from the non-iridescent race (*H. m. rosina, H. e. demophoon*).

### Variation in the distal extension of the red FW band

To further investigate how the collection of phenotypic data altered the loci identified, we phenotyped and mapped individuals for the distal extension of the red FW band using more ‘traditional’ linear measurements (rather than pixel-based pattern extraction). In *H. erato*, the F2 generation had a single QTL on chromosome 13 (LOD = 4.67, p = 0.02), whilst in the BC generation a single significant QTL was found on chromosome 15 (LOD = 5.40, p <0.001) (Figure 3 F, G). No significant QTL were found for *H. melpomene* (Figure 3E). The QTL on chromosomes 13 and 15 included *vvl* and *cortex* respectively within their 95% credible intervals, although the intervals are large (Figure 4, Table 1). The loci explained 19.5% and 23.4% of the variation in the extension of the FW band in the F2 and BC generations, respectively.

In the F2 generation of *H. erato*, the *demophoon* allele for the QTL on chromosome 13 produced a more extended bar and showed slight dominance over the *cyrbia* allele. For the backcross generation, only two genotypes are possible (homozygote *cyrbia* or heterozygote). Surprisingly, individuals with a *cyrbia* genotype at the chromosome 15 locus had longer measures for the distal extension than heterozygotes (Figure 5 C, D).

### Chromosome level analysis to investigate the level of genetic parallelism

Small effect loci are likely to be missed in QTL analyses with small sample sizes due to lack of power. However, using a chromosome level analysis, we can look at the combined effects of multiple small effect loci. For example, if there are many small effect loci on a single linkage group, these may not be detected in the full genome scan but the additive LOD scores could reach significance when comparing effect sizes of whole chromosomes. This can then be used to compare genetic architecture between species. This comparison of maternal alleles provides further evidence for differences in genetics of this trait between species, as there are no parallel significant chromosomes. For *H. erato*, the results are very similar to those found with all markers, although only chromosomes 13 and 18 are detected as significant, with both affecting the multivariate trait, and 18 and 13 affecting the area and linear measurements respectively (Table 1, Figure S4).

For *H. melpomene*, chromosome 20 was found to significantly affect the multivariate trait (LOD = 4.01, p = 0.01), which was not seen when all markers were used, suggesting there may be multiple small effect loci located on this chromosome. LOD scores for both the area and distal extension analyses were very low in *H. melpomene*, but the Z chromosome had the highest LOD score for both, with a significant effect on area. However, most of these offspring are from one cross direction, so sex and maternal Z genotype are highly confounded in this analysis, meaning that this result could reflect sexual dimorphism rather than sex linkage.

To test for parallelism in the combined effect of all markers on each chromosome, we compared the variance explained by each chromosome in *H. erato* and *H. melpomene* and found no correlation between the species in either relative area (r = 0.16, df = 19, p = 0.48) or FW band extension (r = 0.09, df = 19, p = 0.71) (Figure S5). The maternal chromosomes explain little of the overall variation in area, around 15 and 32% in *H. melpomene* and *H. erato* respectively. For FW band extension, 28% of the variation is explained in both species. However, this is not unexpected because the most that these maternal genotypes can explain is 50%, with the rest of the genetic variance coming from the paternal markers.

## Discussion

Our analysis shows that complex variation in the size and shape of red pattern elements, in both *H. melpomene* and *H. erato*, is controlled by a small number of medium to large effect loci. The small amount of phenotypic variation explained by these loci suggests there are likely more loci of minor effect involved. These findings are broadly consistent with the theoretical prediction that an adaptive walk follows a ‘two-step’ process, where single large effect loci evolve initially (during large trait shifts) but smaller modifier loci evolve later to refine the subtle variation (Nicholson 1927; Sheppard et al. 1985; Orr 1998). Multivariate QTL scans for *H. erato* identified a greater number of smaller effect loci in a single scan than those that mapped univariate traits. The inability of univariate measurements to capture the full magnitude or direction of shape changes has been demonstrated by a number of recent studies. Consequently, QTL scans based off such values are likely to be underestimations of the true complexity of the genetic architecture underlying adaptive traits (Maga et al. 2015; Pallares et al. 2016; Rossato et al. 2018).

A major question asked by this study was whether repeated evolution between converging species is reflected by the same or different genetic changes (Parchem et al. 2007; Ng et al. 2008; Nadeau et al. 2014). Although the known major effect locus *optix* was re-used in both species, surprisingly few genetic parallels were found between these two co-mimics. These results suggest that despite parallel patterns of divergence and high levels of gene re-use at major patterning loci, levels of interspecific genetic parallelism are in fact quite low between small effect loci. The transcription factor *optix* already has a well-defined role in the colour fate of scale cells acting as a switch-like regulator between orange/red (ommochrome) and black/grey (melanin) patterns (Baxter et al. 2008; Reed et al. 2011; Nadeau et al. 2014). Our results are consistent with previous QTL studies and recent gene editing experiments, which have found that genetic variation around the *optix* gene can also have subtle effects on the size and shape of red pattern elements (Lewis et al. 2019).

Along with *optix*, other loci control variation in size and shape of these red elements (Nadeau 2016). The most prominent of these is the *WntA* gene on chromosome 10, which has been shown to control the placement of melanic scales in both *H. melpomene* and *H. erato* (Naisbit et al. 2003; Martin et al. 2012; Jiggins 2017). As we specifically investigated shape differences in red FW pattern elements, it is surprising that we did not find any QTL which might contain *WntA*. This result indicates that the shape of red pattern elements studied in the crosses here may follow a divergent process to what has been studied in other crosses previously, and that alleles other than those contained within *WntA* are having an effect here. Alternatively, our QTL analysis may have been unable to detect effects of *WntA* with the sample sizes used here, if they had a small effect (Beavis 1998). The linkage maps for both species had a marker less than 170kbp from the position of *WntA*, so it is likely that any medium to large effect alleles at this locus would have been detected.

In contrast, we do find QTL on chromosome 13 containing the *vvl* gene. This locus is less well studied but has previously been identified as controlling forewing band shape variation in both *H. erato* (Sheppard et al. 1985; Nadeau et al. 2014; Van Belleghem et al. 2017) and *H. melpomene* (Baxter et al. 2009; Morris et al. 2019). Our finding of a QTL for distal extension of the forewing band that overlaps with the gene *cortex* in *H. erato* is more surprising. Often, *cortex* is described as having control over yellow/white (rather than red) wing pattern elements in both species (Joron et al. 2006; Nadeau 2016). However, some studies have begun to acknowledge the potential alternative effects of this locus. Nadeau (2016) suggested that the effect of *cortex* on colour might vary with the developmental stage that it is up-regulated in, whilst others have found evidence for an epistatic interaction between *cortex* and *optix* in *H. melpomene* (Salazar 2012; Martin et al. 2014). Thus, it is possible that *cortex* is controlling red band shape through an epistatic relationship with *optix*, which may explain why it is detected in the backcross but not the intercross F2.

In *H. erato*, two novel patterning loci were identified on chromosomes 11 and 13. These QTL likely represent some of the small effect loci that form the latter half of the hypothesised ‘two-step’ model (Orr 1998). No previous studies have identified a locus controlling forewing red pattern variation on chromosome 11 in either species.

Further support for a lack of genetic parallelism comes from the analysis of only maternal alleles. We do not see evidence of the same chromosomes containing genes for forewing band patterning across species. While the scans for *H. erato* showed the same chromosomes to be significant as the full analysis (13 and 18), scans of *H. melpomene* revealed chromosomes 20 and Z as significant. These were not seen in the full analysis, suggesting there may be multiple small effect loci on these chromosomes. There is previous evidence for a locus controlling forewing band shape on chromosome 20 in *H. melpomene* (Nadeau et al. 2014) which could be a possible candidate.

When phenotypic convergence is reflected by convergent genetic mechanisms it supports the idea that genetic evolution is fairly constrained, and therefore that adaptive evolution may be predictable (Stern and Orgogozo 2008; Losos 2011). Comparative mapping studies in *Heliconius* find evidence for homologous wing patterning loci, thus leading to a general perception that genetic evolution is surprisingly repeatable (Papa et al. 2008; Martin et al. 2014). However, despite finding genetic parallels with *optix* in this study, other QTL identified do not show similarities. In addition, the phenotypic effect of *optix* appears to differ between species, with an effect on band area in *H. erato* but only on band shape in *H. melpomene*. This suggests that evolution at the genetic level in *Heliconius* is not as predictable as first thought. Thus, indicating that perhaps large phenotypic changes, relating to the initial steps of the hypothesised ‘two-step’ model, are much more constrained, and therefore repeatable, than the latter half of the model (relating to subtler trait shifts).

A question that then remains is, what makes certain loci more likely to evolve in parallel over others? It is possible that finding genetic ‘hotspots’ is a mere reflectance of biases in the research; focusing on known candidate genes is considerably easier than identifying novel ones (Conte et al. 2012). Additionally, identifying QTL of large effect is easier than identifying those of small effect (Rockman 2011). One reason that certain loci are highly re-used in adaptation, whilst others are not, could be their genetic architecture. *Optix* has been shown to have a complex modular architecture, consisting of multiple cis-regulatory modules (Reed et al. 2011; Wallbank et al. 2016; Van Belleghem et al. 2017). Pleiotropy, where a gene affects multiple different traits, is often assumed to be purged in evolution as it rarely results in solely advantageous changes. However, genes that are able to integrate multiple upstream regulatory elements and in turn, alter specific phenotypes without causing any knock-on deleterious effects, will be favoured in adaptive evolution (Stern and Orgogozo 2008; Gompel and Prud’homme 2009). Thus, it could be hypothesised that these loci have become ‘hotspots’ for adaptive change across the *Heliconius* genus due to their ability to guide large effect mutations, without deleterious effects on other traits (Reed et al. 2011; Pavličev and Cheverud 2015; Jiggins 2017).

## Conclusion

Previous studies have often focused on major effect loci that affect discrete pattern changes in *Heliconius*. In contrast, this study has demonstrated that, with multivariate trait analysis, QTL mapping can be used to identify a greater number of smaller effect loci in a single analysis, in turn questioning the role of a simple genetic ‘tool-kit’ in *Heliconius*. Combining this study with previous findings builds a picture of a genetic architecture that is consistent with the theoretical ‘two-step’ model of evolution. However, finding that genetic parallels are not as widespread as first expected suggests that the individual path each species takes along this adaptive walk is likely less predictable than previously thought.

## Acknowledgements

We thank the governments of Ecuador and Panama for permission to collect butterflies. Thanks to Darwin Chalá, Emma Curran, Juan López and Gabriela Irazábal for their assistance with the crosses. This work was funded by a NERC Independent Research Fellowship to NJN (NE/K008498/1). MNB was funded by the NERC ACCE DTP (NE/L002450/1).

## Author Contributions

Analyses were run by HEB and MNB. Linkage maps were constructed by PR and MNB. The crosses were performed by PAS, CM, MNB and NJN. The study was devised and co-ordinated by NJN. HEB and MNB wrote the manuscript. All authors read and commented on the manuscript.

## Data Availability

Sequence data has been deposited in the European Nucleotide Archive under project number PRJEB38330. Photographs of all samples will be available at doi.org/10.5281/zenodo.3799188.

