## Supplementary for "Limited genetic parallels underlie convergent evolution of quantitative pattern variation in mimetic butterflies"

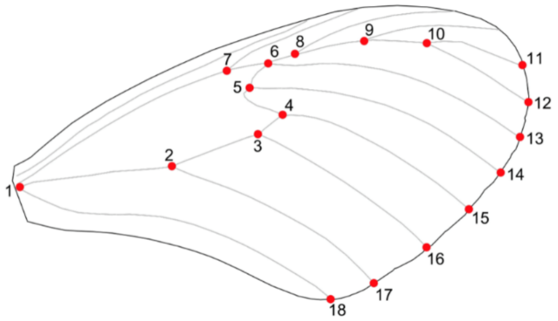


Figure S1. Diagram of the landmarks used to align each image; points must be in the same order for each sample (Source: Van Belleghem, 2017).

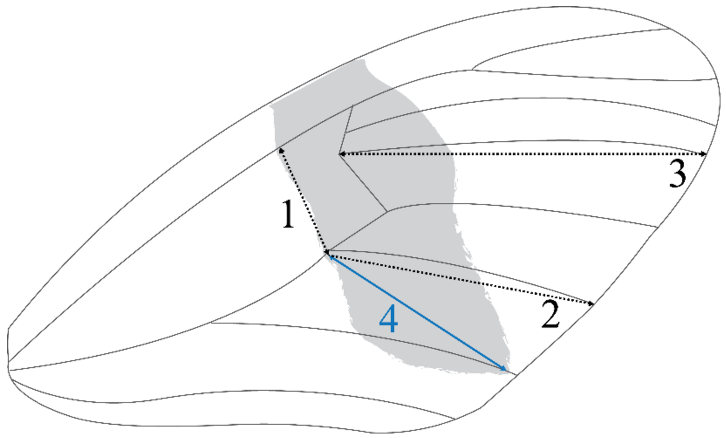


Figure S2. Schematic diagram to represent how the extension of the red FW band was measured. Dashed lines (1, 2, 3) represent measurements taken to standardise for wing size, whilst the full arrow (4) shows where the 'distal extension measurement' was taken (Adapted from: Baxter et al. 2009; Brien et al. 2018).

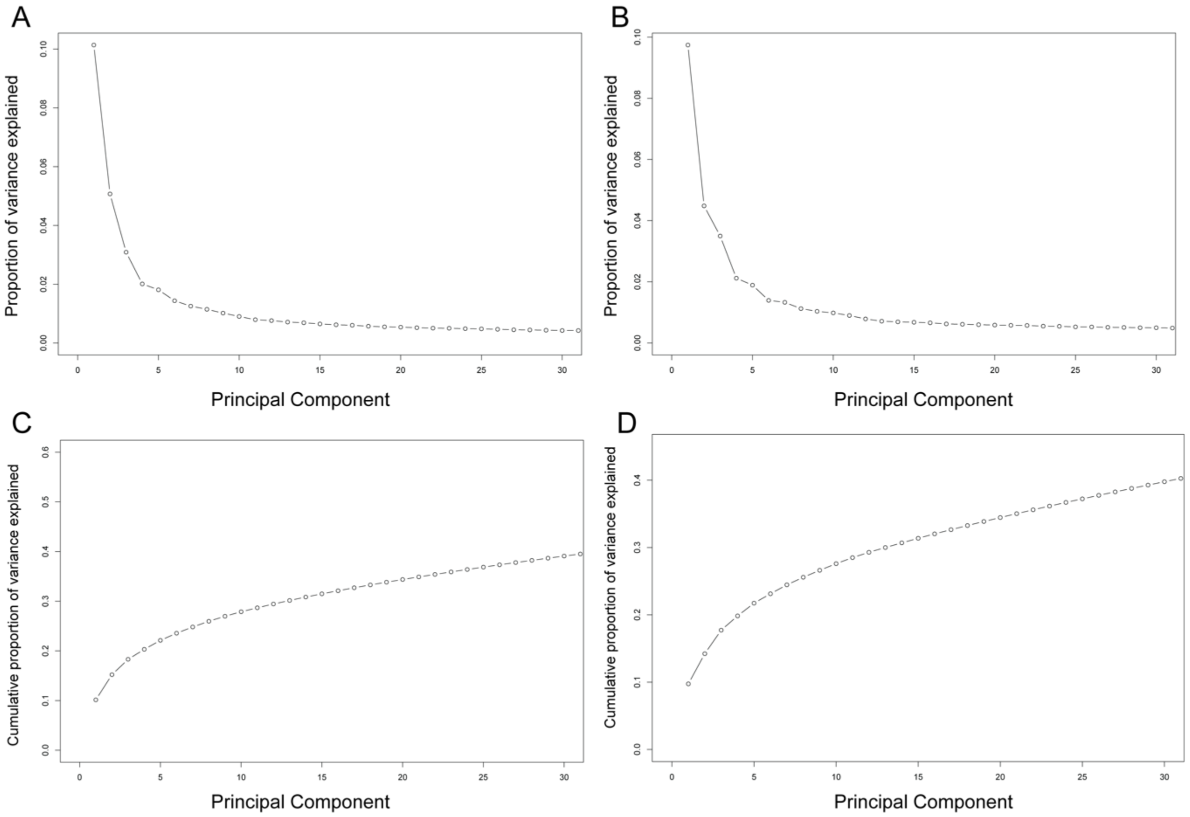


Figure S3. Scree plots showing the proportion of variance explained (A, B) and the cumulative proportion of variance explained (C, D) for each of the first 30 PCs (from PCA analyses across all the generations) for quantitative red forewing variation, conducted on *H. melpomene* (A, C) and *H. erato* (B, D).

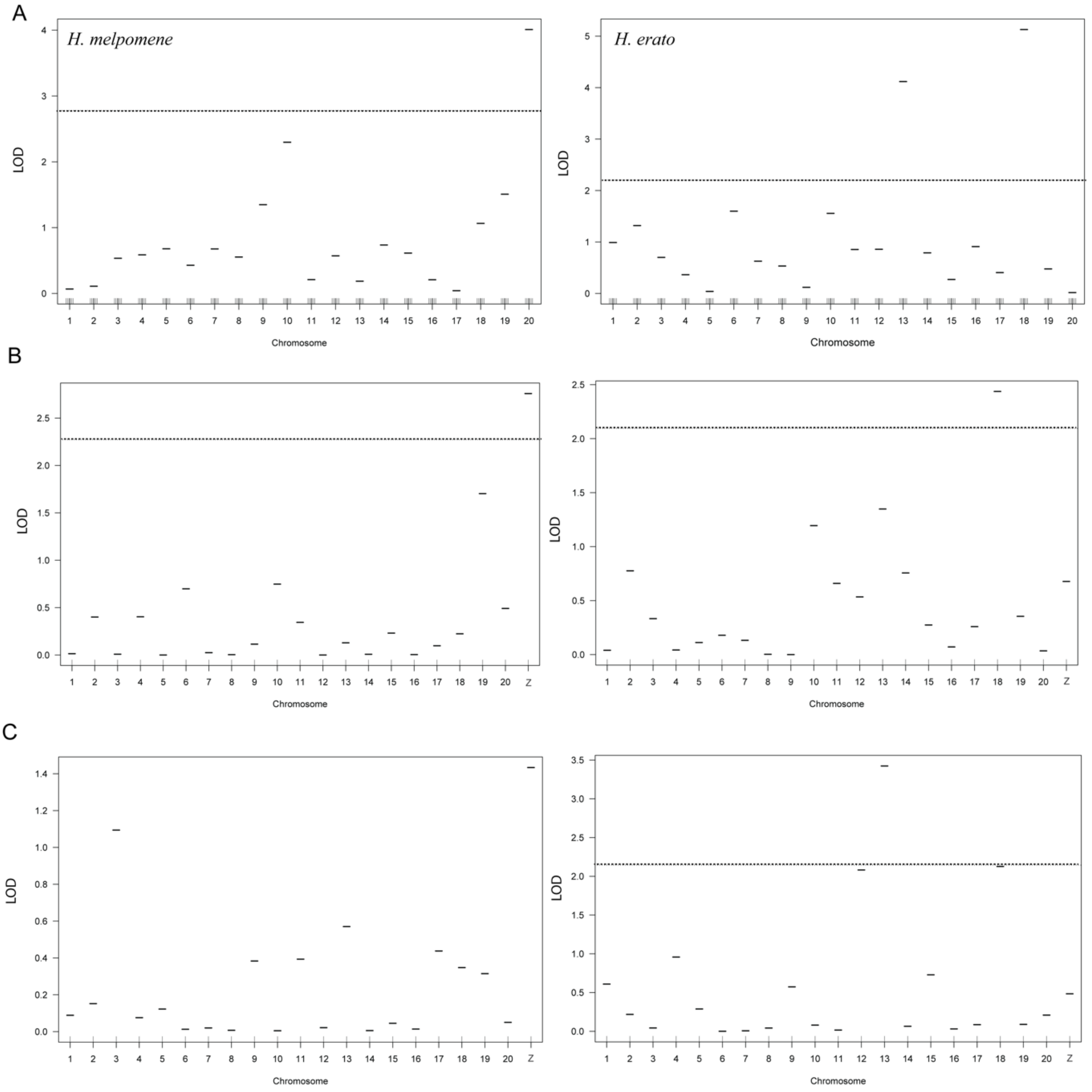


Figure S4. Chromosome level QTL scans in *H. melpomene* (left) and *H. erato* (right) for (A) shape, (B) relative area and (C) FW band extension.

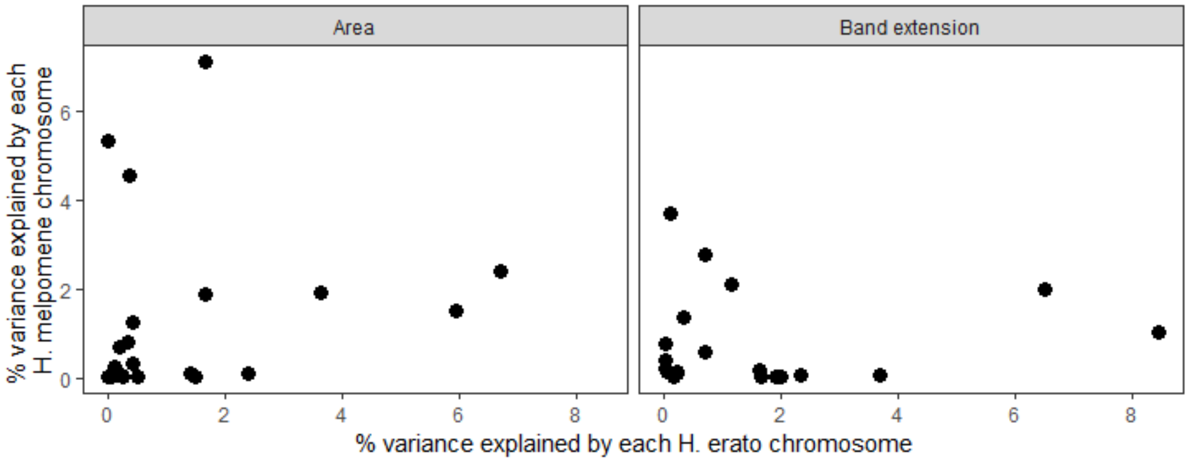


Figure S5. No correlation was found between the percentage of variation in relative area or in FW band extension explained by *H. melpomene* chromosomes and *H. erato* chromosomes.

Table S1: Information on the *Heliconius erato* and *Heliconius melpomene* crosses which produced offspring that were included in the analyses. Parents of F2 crosses and backcrosses have the cross they originated from in brackets.

| **Cross ID** | **Cross type** | **Father ID** | **Mother ID** | **Offspring phenotyped for:** | | **Sequenced to produce linkage maps** |
|---|---|---|---|---|---|---|
| | | | | Shape and area | FW band extension | |
| ***Heliconius erato*** | | | | | | |
| EC01F1 | *demophoon* ♂x *cyrbia* ♀ | 14N012 | 14N011 | 17 | | 4 |
| EC03F1 | *cyrbia* ♂ x *demophoon* ♀ | Unknown | 14N047 | 2 | | |
| EC10F1 | *cyrbia* ♂ x *demophoon* ♀ | 14N065 | 14N064 | 14 | | 3 |
| EC39F1 | *cyrbia* ♂ x *demophoon* ♀ | 14N339 | 14N338 | 7 | | 2 |
| EC45F1 | *demophoon* ♂x *cyrbia* ♀ | 14N359 | 14N358 | 15 | | 2 |
| EC11BC | *cyrbia* ♂ x (*demophoon* ♂ x *cyrbia* ♀) | 14N067 | 14N066 (EC01) | 1 | | |
| EC12BC | *cyrbia* ♂ x (*demophoon* ♂ x *cyrbia* ♀) | 14N069 | 14N068 (EC01) | 13 | | |
| EC41BC | *cyrbia* ♀ x (*cyrbia* ♂ x *demophoon* ♀) | 14N348 (EC10) | 14N347 | 50 | 40 | 40 |
| EC13F2 | *demophoon* maternal grandfather | 14N078 (EC01) | 14N077 (EC01) | 5 | 3 | 3 |
| EC15F2 | *demophoon* maternal grandfather | 14N093 (EC01) | 14N089 (EC01) | 6 | 5 | 5 |
| EC17F2 | *cyrbia* maternal grandfather | 14N112 (EC01) | 14N111 (EC10) | 60 | 56 | 56 |
| EC18F2 | *cyrbia* maternal grandfather | 14N114 (EC01) | 14N113 (EC10) | 16 | 14 | 14 |
| EC53F2 | *cyrbia* maternal grandfather | 14N396 (EC39) | 14N395 (EC39) | 23 | 21 | 21 |
| EC56F2 | *demophoon* maternal grandfather | 14N428 (EC39) | 14N427 (EC45) | 3 | | |
| **Total** | | | | 232 | 139 | 150 |

| | | | | | | |
|---|---|---|---|---|---|---|
| ***Heliconius melpomene*** | | | | | | |
| EC05F1 | *cythera* ♂x *rosina* ♀ | 14N052 | 14N051 | 13 | | |
| EC07F1 | *cythera* ♂x *rosina* ♀ | 14N059 | 14N058 | 11 | | |
| EC09F1 | *rosina* ♂ x *cythera* ♀ | 14N063 | 14N062 | 20 | | |
| EC26F1 | *cythera* ♂x *rosina* ♀ | 14N219 | 14N220 | 4 | | |
| EC27F1 | *rosina* ♂ x *cythera* ♀ | 14N221 | 14N222 | 12 | | |
| EC35F1 | *cythera* ♂x *rosina* ♀ | Unknown | 14N315 | 2 | | |
| EC38F1 | *rosina* ♂ x *cythera* ♀ | 14N337 | 14N336 | 5 | | |
| EC48F1 | *rosina* ♂ x *cythera* ♀ | 14N366 | Unknown | 16 | | 2 |
| EC49F1 | *cythera* ♂x *rosina* ♀ | 14N368 | 14N367 | 13 | | 3 |
| EC51F2 | *rosina* maternal grandfather | 14N379 (EC35) | 14N385 (EC38) | 52 | | |
| EC57F2 | *rosina* maternal grandfather | 14N452 (EC38) | 14N451 (EC48) | 30 | | |
| EC63F2 | *cythera* maternal grandfather | 14N480 (EC48) | 14N479 (EC49) | 59 | 52 | 52 |
| EC65F2 | *cythera* maternal grandfather | 14N563 (EC48) | 14N562 (EC49) | 52 | 54 | 54 |
| EC69F2 | *rosina* maternal grandfather | 14N605 (EC49) | 14N604 (EC48) | 8 | 7 | 7 |
| **Total** | | | | 297 | 113 | 118 |

Table S2: Genetic map summary for *H. erato*. The *H. erato* genome is 380Mb meaning a centimorgan (cM) in this map has a physical distance of 330Kb on average.

| **Linkage group** | **Number of markers** | **Length (cM)** | **Average spacing (cM)** | **Maximum spacing (cM)** |
|---|---|---|---|---|
| **1** | 303 | 56.3 | 0.2 | 3.9 |
| **2** | 247 | 42.9 | 0.2 | 4.7 |
| **3** | 219 | 62.8 | 0.3 | 7.8 |
| **4** | 438 | 56.7 | 0.1 | 6.8 |
| **5** | 371 | 41.9 | 0.1 | 2.3 |
| **6** | 333 | 65.0 | 0.2 | 9.0 |
| **7** | 228 | 51.4 | 0.2 | 5.5 |
| **8** | 333 | 43.4 | 0.1 | 6.4 |
| **9** | 208 | 52.9 | 0.3 | 6.4 |
| **10** | 373 | 49.9 | 0.1 | 4.5 |
| **11** | 265 | 75.7 | 0.3 | 7.2 |
| **12** | 340 | 57.0 | 0.2 | 3.9 |
| **13** | 244 | 50.1 | 0.2 | 3.9 |
| **14** | 180 | 60.2 | 0.3 | 6.7 |
| **15** | 335 | 69.0 | 0.2 | 7.2 |
| **16** | 229 | 54.0 | 0.2 | 3.1 |
| **17** | 199 | 69.1 | 0.3 | 11.7 |
| **18** | 234 | 52.7 | 0.2 | 4.7 |
| **19** | 299 | 54.4 | 0.2 | 5.3 |
| **20** | 160 | 46.1 | 0.3 | 3.1 |
| **Z** | 110 | 50.9 | 0.5 | 10.8 |
| **Overall** | **5648** | **1162.4** | **0.2** | **11.7** |

Table S3: The genetic linkage map used for the *H. melpomene* analysis contained 2163 markers across 21 linkage groups.

| **Linkage group** | **Number of markers** | **Length (cM)** | **Average spacing (cM)** | **Maximum spacing (cM)** |
|---|---|---|---|---|
| **1** | 123 | 69.9 | 0.6 | 2.8 |
| **2** | 103 | 65.0 | 0.6 | 2.8 |
| **3** | 97 | 65.6 | 0.7 | 4.2 |
| **4** | 100 | 72.1 | 0.7 | 4.4 |
| **5** | 99 | 79.1 | 0.8 | 6.0 |
| **6** | 108 | 71.2 | 0.7 | 5.8 |
| **7** | 100 | 66.6 | 0.7 | 5.3 |
| **8** | 90 | 72.6 | 0.8 | 5.2 |
| **9** | 87 | 59.7 | 0.7 | 3.9 |
| **10** | 134 | 82.5 | 0.6 | 2.9 |
| **11** | 90 | 68.0 | 0.8 | 4.8 |
| **12** | 107 | 63.1 | 0.6 | 1.9 |
| **13** | 122 | 74.8 | 0.6 | 6.5 |
| **14** | 89 | 65.3 | 0.7 | 5.1 |
| **15** | 102 | 61.7 | 0.6 | 2.3 |
| **16** | 99 | 72.7 | 0.7 | 4.6 |
| **17** | 107 | 75.7 | 0.7 | 5.0 |
| **18** | 125 | 76.5 | 0.6 | 2.8 |
| **19** | 119 | 72.9 | 0.6 | 2.5 |
| **20** | 111 | 73.9 | 0.7 | 4.4 |
| **Z** | 51 | 61.0 | 1.2 | 5.3 |
| **Overall** | 2163 | 1469.9 | 0.7 | 6.5 |

Table S4: Locations of the ‘toolkit’ loci in the respective reference genomes (scaffold name and location in bp) (Source: Lepbase).

| Gene | *H. melpomene (v2.5)* | *H. erato (v1)* |
|---|---|---|
| *Optix* | Hmel218003o:705,604-706,407 | Herato1801:1,239,943-1,251,211 |
| *Cortex* | Hmel215003o:1,413,776-1,533,113 | Herato1505:2,074,108-2,087,841 |
| *Vvl* | Hmel213001o:10,310,374-10,311,495 | Herato1301:14,341,251–14,412,364 |
